# Towards fast and reliable simultaneous EEG-fMRI analysis of epilepsy with automatic spike detection

**DOI:** 10.1101/361113

**Authors:** Amir Omidvamia, Magdalena A. Kowalczyk, Mangor Pedersen, Graeme D. Jackson

**Affiliations:** The Florey Institute of Neuroscience and Mental Health, Austin Campus, Melbourne, VIC, Australia; The University of Melbourne, VIC, Australia; Department of Neurology, Austin Health, Melbourne, VIC, Australia

**Keywords:** EEG, fMRI, focal epilepsy, spike detection, interictal discharge, matched filtering

## Abstract

**Objective:** The process of manually marking up epileptic spikes for simultaneous electroencephalogram (EEG) and resting state functional MRI (rsfMRI) analysis in epilepsy studies is a tedious and subjective task for a human expert. The aim of this study was to evaluate whether automatic EEG spike detection can facilitate EEG-rsfMRI analysis, and to assess its potential as a clinical tool in epilepsy.

**Methods:** We implemented a fast algorithm for detection of uniform interictal epileptiform discharges (IEDs) in one-hour scalp EEG recordings of 19 refractory focal epilepsy datasets (from 16 patients) who underwent a simultaneous EEG-rsfMRI recording. Our method was based on *matched filtering* of an IED template (derived from human markup) used to automatically detect other ‘similar’ EEG events. We comprehensively compared simultaneous EEG-rsfMRI results between automatic IED detection and standard analysis with human EEG markup only.

**Results:** In contrast to human markup, automatic IED detection takes a much shorter time to detect IEDs and export an output text file containing spike timings. In 13/19 focal epilepsy cases, statistical EEG-rsfMRI maps based on automatic spike detection method were comparable with human markup, and in 6/19 focal epilepsy cases it revealed additional brain regions not seen with human EEG markup. Additional events detected by our automated method independently revealed similar patterns of activation to a human markup. Overall, automatic IED detection provides greater statistical power in EEG-rsfMRI analysis compared to human markup in a short timeframe.

**Conclusions:** Automatic spike detection is a simple and fast method that *can* reproduce comparable and, in some cases, even superior results compared to the common practice of manual EEG markup in EEG-rsfMRI analysis of epilepsy.

**Significance:** Our study shows that IED detection algorithms can be effectively used in epilepsy clinical settings. This work further helps in translating EEG-rsfMRI research into a fast, reliable and easy-to-use clinical tool for epileptologists. Our IED detection approach will be publicly available as a MATLAB package at: *https://github.com/omidvarnia/Automatic_focal_spike_detection*.

**Highlights:**

- Automatic spike detection increases the number of detected uniform epileptic interictal discharges and enhances statistical power of EEG-rsfMRI inter-subject variability maps,
- Automatic spike detection can identify additional activated brain regions with presumed epileptogenic focus not seen in standard analysis based on human markup,
- Automatic spike detection can shorten the IED identification process.

## 1. Introduction

The ability to simultaneously acquire electroencephalogram (EEG) and functional MRI (fMRI) is clinically valuable in epilepsy, where EEG is used to detect the times at which epileptiform activity is present (due to its *high temporal resolution*) and fMRI is used to map the spatial location of the associated EEG activity (due to its *high spatial resolution*) (Allen et al. 1998). Surface EEG signals are associated with postsynaptic cortical currents of large pyramidal neurons, which are perpendicularly aligned to the cortical surface (Nunez and Srinivasan 2006; Daunizeau et al. 2010). FMRI data analysis, on the other hand, relies on the blood oxygen level-dependent (BOLD) contrast of active/inactive brain (Ogawa et al. 1990). Specifically, interictal epileptiform discharges (IEDs) are visible on scalp EEG when synchronous neuronal firing occurs in a substantial portion of cortex (between 10 and 20 *cm*^2^’) (Tao et al. 2007). The significant prognostic value of IEDs for patients with newly diagnosed seizure disorders is well established as they have the potential of revealing a patient’s epileptogenic focus, even in cases with no other imaging evidence (Wirrell 2010).

In order to extract the IED-induced BOLD changes in simultaneous EEG-rsfMRI recordings, a human expert inspects long EEG recordings using a ‘mental template’ for similar epileptogenic events in newly observed EEG signals. The accuracy of human markup is affected by several factors: *i*) it is a subjective, time-consuming and cumbersome procedure which is prone to the risk of missing spikes due to fatigue, in particular in long recordings, *ii*) it needs extensive experience and training, thus being expensive in clinical settings, *iii*) the statistical power of resulted epileptogenic networks may be significantly reduced by markup inconsistency (Waites et al. 2005; Flanagan et al. 2009). It is, therefore, important to obtain more objective and rapid ways of marking epileptogenic spikes in scalp EEG, as this step limits its clinical usefulness and reliability in routine clinical centers.

Several attempts have been made to mitigate the burden of IED markup by a human expert in epilepsy studies (Grouiller et al. 2011; Geerts 2012; Tousseyn et al. 2014; Scheuer et al. 2017). In this context, there is a critical question regarding all existing techniques: can automated detection of scalp-level interictal epileptogenic discharges replicate, and possibly improve the performance of human markup? In the current study, we implemented a fast IED detection algorithm for scalp interictal EEG recordings with roughly uniform IEDs based on matched filtering. In this detection strategy, a known signal, or *template,* is searched throughout an unknown signal in order to find similar events. Specifically, we compared statistically significant BOLD changes across the brain obtained by standard EEG-rsfMRI analysis, based on manually marked spikes by an expert as well as detected IEDs by the algorithm. Our aim was two-fold: i) evaluate if automatic spike detection can improve performance of EEG-rsfMRI analysis in epilepsy studies, and ii) evaluate whether it has the potential to become a user-friendly and efficient tool for EEG experts.

## 2 Methods

### 2.1 Subjects and EEG-rsfMRI acquisition

Since 2012, 35 patients with refractory focal epilepsy underwent an EEG-rsfMRI study as a part of pre-surgical work-up through the Austin Health Comprehensive Epilepsy Program. In total, 19 patients were excluded from this study: 13 patients had no active EEG during EEG-rsfMRI and four patients had significant scanner and/or EEG artefacts. Two patients had ictal, but no IEDs during the scan. We subsequently analyzed simultaneous EEG-rsfMRI data of the remaining 16 patients with focal epilepsy (aged 11-60, 8 females - see Table 1 for more details). All patients had one hour of continuous concurrent EEG-rsfMRI recording. Patients were included in the study if they had relatively frequent interictal epileptiform discharges (>15 IEDs/hour) during the scan. The study was approved by the Austin Health Human Research Ethics Committee and all patients gave written consent to participate in the study.

**Table 1:** Patient details and clinical information

| Patient | Age | Sex | Age of seizure onset | Epilepsy type | MRI finding | PET finding | Ictal SPECT finding | VEM finding |
| --- | --- | --- | --- | --- | --- | --- | --- | --- |
| 1 | 41y | M | 10y | Left TLE | Left hippocampal sclerosis, multiple tubers in the left temporal lobe, left parietal region and left anterior cingulate | Left temporal hypometabolism | Left temporal hyperperfusion | IEDs: left temporal with rare discharges on there right; Ictal: left hemisphere |
| 2 | 42y | F | 7y | Left TLE | Periventricular nodular heterotopia, more severe on left | NA | NA | IEDs and ictal: left temporal |
| 3 | 29y | F | 16y | Left TLE | MRI-negative | Left anterior temporal hypometabolism | Left temporal hyperperfusion | IEDs: left temporal |
| 4 | 39y | F | 3mo | Right FLE or multifocal | MRI-negative | Right frontal, right temporal hypometabolism | Right superior frontal hyperperfusion | IEDs and ictal: right frontal |
| 5 | 40y | M | 16y | Right TLE | Lesion negative, previous right hippocampal sclerosis removal | Right temporal hypometabolism, left temporal hypermetabolism | Right frontal and posterior temporal hyperperfusion | IEDs: bitemporal and right posterior temporal; Ictal: bilateral temporal |
| 6 | 26y | M | 13y | Right FLE | MRI-negative | Left frontal hypometabolism | Right temporal, mesial frontal and right frontal polar cortex hyperperfusion | IEDs and ictal: right central and right frontal |
| 7 | 23y | M | 16y | Right hemisphere focal | Right periventricular nodular heterotopia, right temporal peri-sylvian and parietal polymicrogyria, regions of cortical dysplasia | Right polar, anteromesial and lateral temporal hypometabolism | Right temporal hyperperfusion | IEDs: right fronto-temporal; Ictal: temporal |
| 8 | 41y | F | 16y | Left TLE | MRI-negative | No definitive hypometabolism | Left temporal hyperperfusion | IEDs: left temporal; Ictal: right hemisphere and left posterior quadrant |
| 9 | 26y | F | 5y | Multifocal | Bilateral periventricular nodular heterotopia, ectopic posterior pituitary, extensive left cerebellar hypoplasia, left posterior schizencephaly with polymicrogyria | Left posterior lateral temporal hypometabolism. | Right parieto-temporal-occipital hyperperfusion | IEDs: bilateral fronto-central, left posterior quadrant; Ictal: bilateral |
| 10 | 14y | M | 12y | Right FLE or multifocal | MRI-negative | Right parietal and frontal hypometabolism | Right parietal and frontal hypoperfusion | IEDs: right frontal and fronto-central |
| 11 | 19y | F | 12y | Right parietal | Right postcentral bottom of sulcus dysplasia | Right postcentral hypometabolism | Right postcentral hyperperfusion | IEDs: right centro-temporal |
| 12 | 11y | F | 10y | Left PHG or FLE | Left PHG dysplasia, particularly posteriorly | NA | NA | IEDs: left frontal, rare right frontal; Ictal: left frontal |
| 13 | 27y | F | 8y | Right TLE | MRI-negative | Right posterior temporal hypometabolism | Right posterior temporal hyperperfusion | IEDs: right posterior inferior temporal; Ictal: right temporal. |
| 14 | 25y | M | 11y | Right FLE | Abnormality in the inferior bank of the calcarine sulcus around the right fusiform gyrus | Right frontal hypermetabolism | Right frontal hyperperfusion | IEDs: right anterior quadrant; Ictal: right frontal and temporal |
| 15 | 53y | M | 16y | Left FLE | Left superior-frontal dysplasia, previous left superior frontal resection | Left mesial inferior frontal hypermetabolism, at the posteroinferior margin of the resection cavity | Left frontal hyperperfusion | IEDs and ictal: left frontal |
| 16 | 60y | M | 26y | Right TLE | Right insular surgical defect, previous right inferior frontal resection | Right anterior temporal hypometabolism | Right anterior temporal hyperperfusion with surrounding hypoperfusion | IEDs: right temporal |

Abbreviations: M = male, F = female, y = year, mo = month, NA = not available, TLE = temporal lobe epilepsy, FLE = frontal lobe epilepsy, PHG = parahippocampal gyrus, MRI = magnetic resonance imaging, PET = positron emission tomography, SPECT = single photon emission computed tomography, VEM = video electroencephalogram monitoring, IEDs = interictal epileptiform discharges

Each patient was scanned using a 3 Tesla Siemens Skyra system (Erlangen, Germany) with eyes closed and instructed to fall asleep to maximize the probability of detecting epileptogenic events on scalp EEG. FMRI data were acquired using an echo-planar imaging sequence with 44 interleaved 3 mm slices, repetition time = 3 s, echo time = 30 ms, flip angle = 85°, voxel size = 3×3×3 mm, field of view = 216 mm and acquisition matrix size = 72×72. A total of 1200 volumes (60 minutes) of rsfMRI data were used for all patients. A Tļ-weighted anatomical image at 1.2×1.2×1.2 mm resolution was also acquired during each recording session.

Simultaneous EEG data were acquired using a 32-channel MR-compatible EEG cap (BrainCap MR, EasyCap GmbH, Breitbrunn, Germany) according to the 10-20 standard system using a BrainAmp recorder (Brain Products GmbH, Munich, Germany). Data were recorded based on the referential montage at the sampling rate of 5000 Hz with reference to the FCz electrode, but further converted to average reference, and grounded to AFz. Additional channels included echocardiogram and signals from three motion coils that were developed in our lab (Masterton et al. 2007). Sharp spikes and slow-spike-and-waves were the most frequent types of IEDs observed throughout the EEG recordings (see Table 2). Sharp spikes were defined as focal changes of EEG activity in the time domain with duration less than 70 ms. Slow-spike-and-waves were defined as a slow decaying envelope followed by a few fluctuations of EEG activity with a duration less than 200 ms.

**Table 2:**
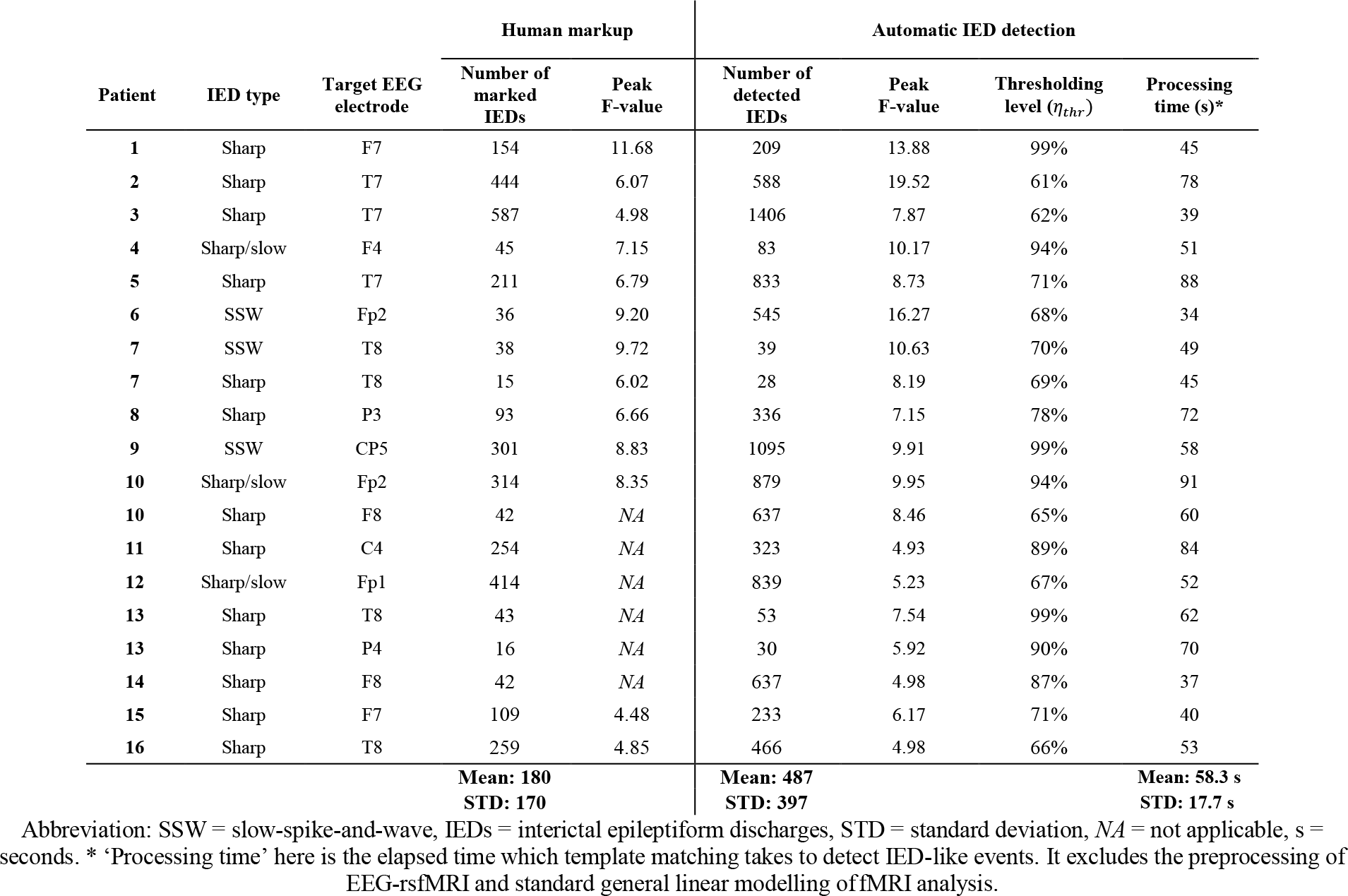
IED identification details for each patient. Here, ‘Peak F-value’ refers to the highest F-value observed in each statistical parametric map of our EEG-rsfMRI analysis.

### 2.2 EEG-rsfMRI preprocessing

All rsfMRI data were preprocessed in MATLAB (MathWorks Inc., Natick, Massachusetts, United States) using Statistical Parametric Mapping toolbox (SPM12, Welcome Department of Imaging Neuroscience, Institute of Neurology, London) and with the aid of the in-house developed iBrain Analysis Toolbox for Statistical Parametric Mapping (available at: *www.brain.org.au/software*)(Abbott et al., 2011). FMRI preprocessing steps included: slice timing correction and re-alignment of the fMRI images for head motion, segmentation of T_1_-weighted images into white matter, grey matter and cerebrospinal fluid areas and coregistration of fMRI images to the patient’s own Tj_-weighted images. Slice time corrected and spatially normalized datasets were smoothed using a spatial Gaussian filter with full-width at half maximum of 8 mm. Also, nuisance signals (6 motion parameters as well as mean white matter and cerebrospinal fluid signals) were regressed out from the data. Slow signal drifts with a period longer than 128 seconds (i.e., frequencies below ~0.008 Hz) were also removed from fMRI time series.

EEG signals were preprocessed using the BrainVision Analyzer software (version 2.0, Brain Products) and the EEGLAB toolbox (Delorme and Makeig 2004). Cardioballistic artifacts were corrected automatically using information obtained via motion artefact detection loops (Masterton et al. 2007; Abbott et al. 2015). Gradient-switching artifacts were corrected by subtracting the averaged scanner artifact template from the continuous EEG recordings using the information of scanner markups (Allen et al. 2000). The preprocessed EEG signals were further downsampled to 250 Hz.

### 2.3 Automatic IED detection using matched filtering

Detection of IEDs in contrast to other non-spike events such as background noise and non-epileptic biological changes can be very challenging, as they may emerge in a wide range of temporal morphologies. In this study, we focused on two types of IEDs with consistent shape in the time domain (i.e., sharp spikes and slow-spike-and-waves) as two of the most widely seen spike types in focal epilepsy. With some simplifying assumptions, the challenge of IED detection can be reframed as a classical detection problem in signal processing where an event of interest is going to be detected throughout a noisy signal. An optimal solution to this problem is offered by the *matched filter theorem* (Geerts 2012; Boashash and Azemi 2014).

Suppose the interictal EEG signal *x*(*t*) of length *T* at a target EEG electrode has been generated through a linear time-invariant system with impulse response function *h*(*t*) and consists of a set of deterministic and real-valued events *s*(*t*) (here, IEDs) plus additive white Gaussian noise *n*(*t*). Two statistical hypotheses can be applied to *x*(*t*):

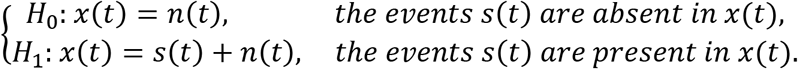

The target EEG electrode is selected where the dominant IED events occur (i.e., the one with greatest mean IED amplitude over all electrodes – see Figure 2-A and Figure 3-A for two examples). The basic idea of matched filtering is to magnify the events of interest in *x*(*t*) by passing it through a linear and shift-invariant filter which is matched to *s*(*t*). It is mathematically shown that the filter *h*(*t*) = *s*(−*t*) can do this job by maximizing the signal-to-noise ratio between *x*(*t*) and *n*(*t*). A test statistic *η*(*t*) is then obtained by filtering *x*(*t*) and sampling the resulted signal at *t* = 0:

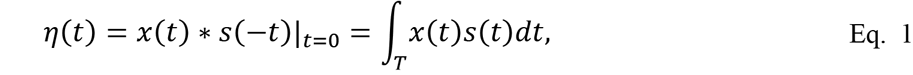

where * denotes the convolution operator. In the case of automatic IED detection, the template *s*(*t*) is defined for each patient by averaging their manually marked epileptogenic events throughout the EEG recorded inside the MRI scanner. It is important to note that matched filtering works best for IEDs with consistent morphology, as they can be represented by a template. In fact, Eq. 1 describes a correlation procedure in the time domain where the ‘similar’ parts of interictal EEG recording to the IED template *s*(*t*) are being highlighted by the matched filter *h*(*t*). The test statistic *η*(*t*) is further normalized by its Euclidean norm and compared with a fixed threshold *η_thr_* for hypothesis testing:

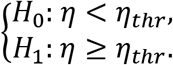

True detection is when we reject *H*_0_ and the event is present. False detection is when we reject *H*_0_, while the event is not present.

### 2.4 Finding the optimal threshold between matching template and new spikes

In order to estimate the optimal threshold *η_thr_* for each patient (i.e., the threshold which can lead to maximum similarity between the template and detected events), we performed a receiver operating characteristic (ROC) analysis. We increased the thresholding level *η_thr_* from 1% to 99% with 1% incremental steps. At each level, we treated the suprathreshold amplitudes as ‘detected IEDs’ and assigned a binary mask to their timing (i.e., 1 for above-threshold and 0 for below-threshold time points). We considered this binary timing as a score set returned by a binary IED classifier whose true class labels were the binary timing of human markup. This resulted in a ROC curve for each thresholding level (99 ROCs in total for each patient). Finally, we chose the optimal threshold as the one associated with the highest area under the ROC curve. Table 2 summarizes the optimum thresholds as well as the results of automatic IED detection in contrast to human markup for all patients.

### 2.5 Simultaneous EEG-rsfMRI analysis based on automatic IED detection and human markup

We used general linear modelling to quantify joint information of IEDs (either automatically detected or manually marked) and simultaneous rsfMRI (Bénar et al. 2002). This process treats IEDs as a binary sequence of delta functions, each of which represents an epileptogenic discharge. We convolved this binary sequence with a canonical hemodynamic response function peaking at 6s to resemble the delay between neural responses and their associated hemodynamic changes. We introduced EEG-derived information to the general linear modelling design matrix as regressors of interest. We identified the spatial epileptic networks associated with IEDs by applying a statistical F-test on the fitted general linear model parameters. We thresholded the statistical parametric maps at a voxel-level p-value of <0.001 and a cluster-level p-value of <0.05 using Gaussian Random Field Theory (Friston et al. 1994).

### 2.6 Comparison of the automatic approach and human markup

The automatic IED detection method was compared with human markup in two ways: (1) based on visual comparison of statistical EEG-rsfMRI parametric maps by two unbiased and experienced epileptologists and EEG experts, and (2) by comparing the processing time used to identify, mark and export timing files of IEDs from EEG recordings using both methods. Figure 1 illustrates the block diagram of this study for a typical dataset.

**Figure 1:**
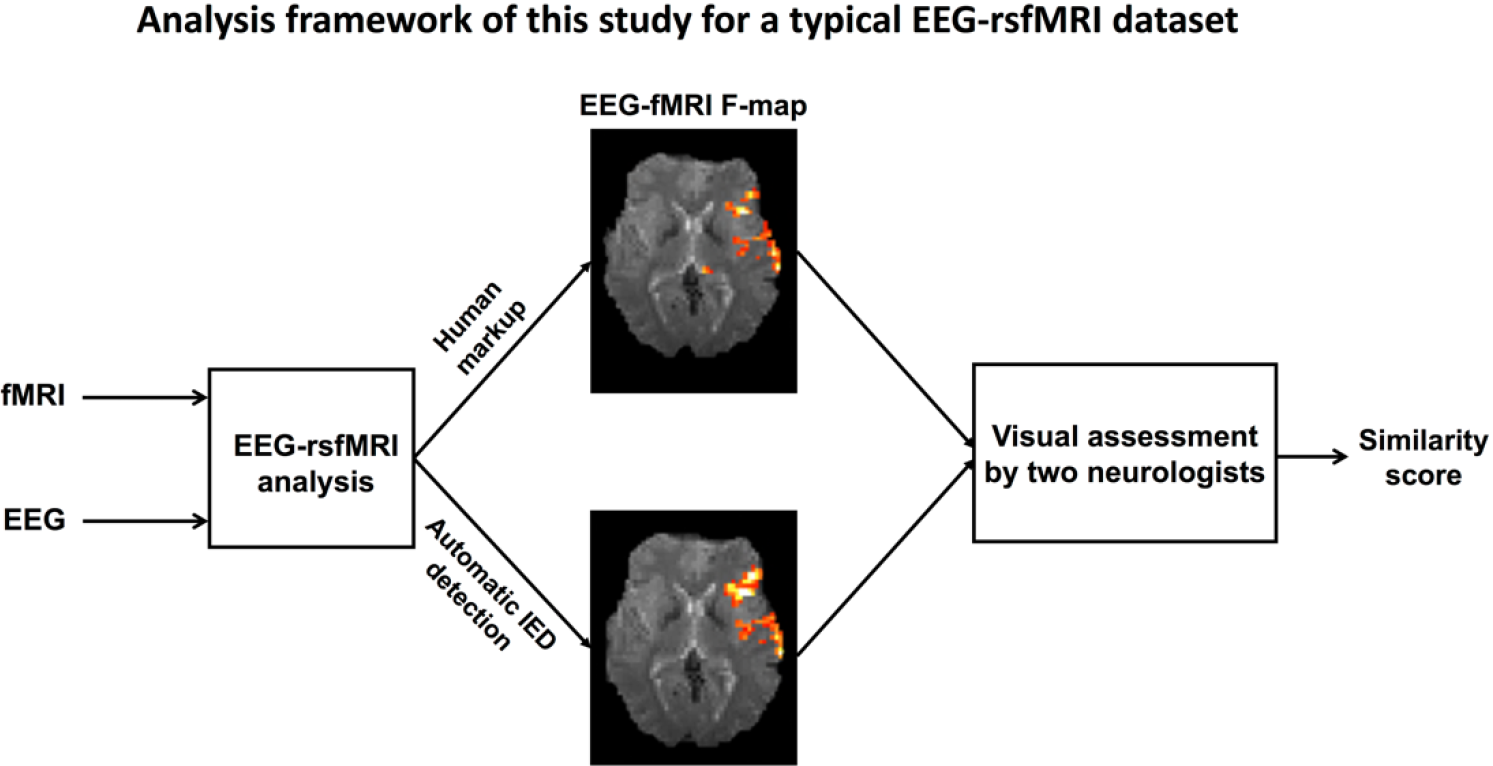
Comparison of automatic IED detection and manual IED markup methods. For each subject, EEG-rsfMRI maps of the two IED markup approaches were given to two independent experts for visual comparison.

#### 2.6.2. Visual evaluation of maps by EEG experts

Spatial EEG-rsfMRI maps (multiple axial slices through the brain in own-space anatomy) for each patient were visually inspected by two independent epileptologists. For each pair of EEG-rsfMRI maps (one based on human markup and the other based on automatic spike detection method - see Figure 1 for an example of this), the two experts were asked to assess the level of visual similarity of maps on a scale from 1 (completely different maps) to 10 (identical maps). If no EEG-rsfMRI clusters were observed in the human markup results, the two EEG experts assessed whether the additional spatial information provided by the automated IED method were clinically informative. Specifically, the experts checked whether the results were in line with available clinical information and other imaging modalities such as PET and ictal SPECT (Table 1).

The experts underwent no specific training or preparation for the study, they were not given any guidance as to how to put their similarity scores and they were asked to not discuss the assessment with each other. The order of presented pairs of F-maps was randomized to avoid any underlying confounding patterns and was kept the same for both experts. Table 3 shows the questionnaire as well as the associated scores. Note that two general classes of IEDs with relatively persistent morphology over time (i.e., slow-spike-and-wave and sharp discharges) were present in our cohort of 16 patients (Table 2). All EEG datasets had one IED type, except for dataset 7, 10 and 13 having two types. Consequently, 19 different IED events from 16 patients were included in this study. Figure 2 and Figure 3 illustrate the results of two exemplary datasets (Patients 1 and 6), while the complete set of results can be found in Figure 4 and Figure 5.

**Table 3:** The questionnaire which was used by two independent EEG experts to score the similarity between the EEG-rsfMRI statistical maps associated with automatically detected IEDs and human markup.

| In general, does the automated IED detection method provide comparable results with the standard human markup method? YES/NO. |  |  |
| --- | --- | --- |
| On a scale from 1 (completely different) to 10 (identical), how similar are these two maps in your opinion? |  |  |
| Patient no. | Expert 1 | Expert 2 |
| 1 | Yes, 9 | Yes, 9 |
| 2 | Yes, 7 | Yes, 7 |
| 3 | No, 4 | Yes, 4 |
| 4 | Yes, 6 | Yes, 6 |
| 5 | Yes, 7 | Yes, 7 |
| 6 | Yes, 4 | Yes, 6 |
| 7 (sharp) | Yes, 4 | Yes, 6 |
| 7 (SSW) | Yes, 9 | Yes, 8 |
| 8 | Yes, 4 | Yes, 4 |
| 9 | Yes, 6 | Yes, 7 |
| 10 (sharp slow) | Yes, 9 | Yes, 9 |
| 10 (sharp) | N/A | N/A |
| 11 | N/A | N/A |
| 12 | N/A | N/A |
| 13 (sharp T8) | N/A | N/A |
| 13 (sharp P4) | N/A | N/A |
| 14 | N/A | N/A |
| 15 | No, 2 | No, 3 |
| 16 | No, 4 | No, 5 |
| Mean | 6.1 | 6.5 |
| Standard deviation | 2.4 | 1.9 |

**Figure 2:**
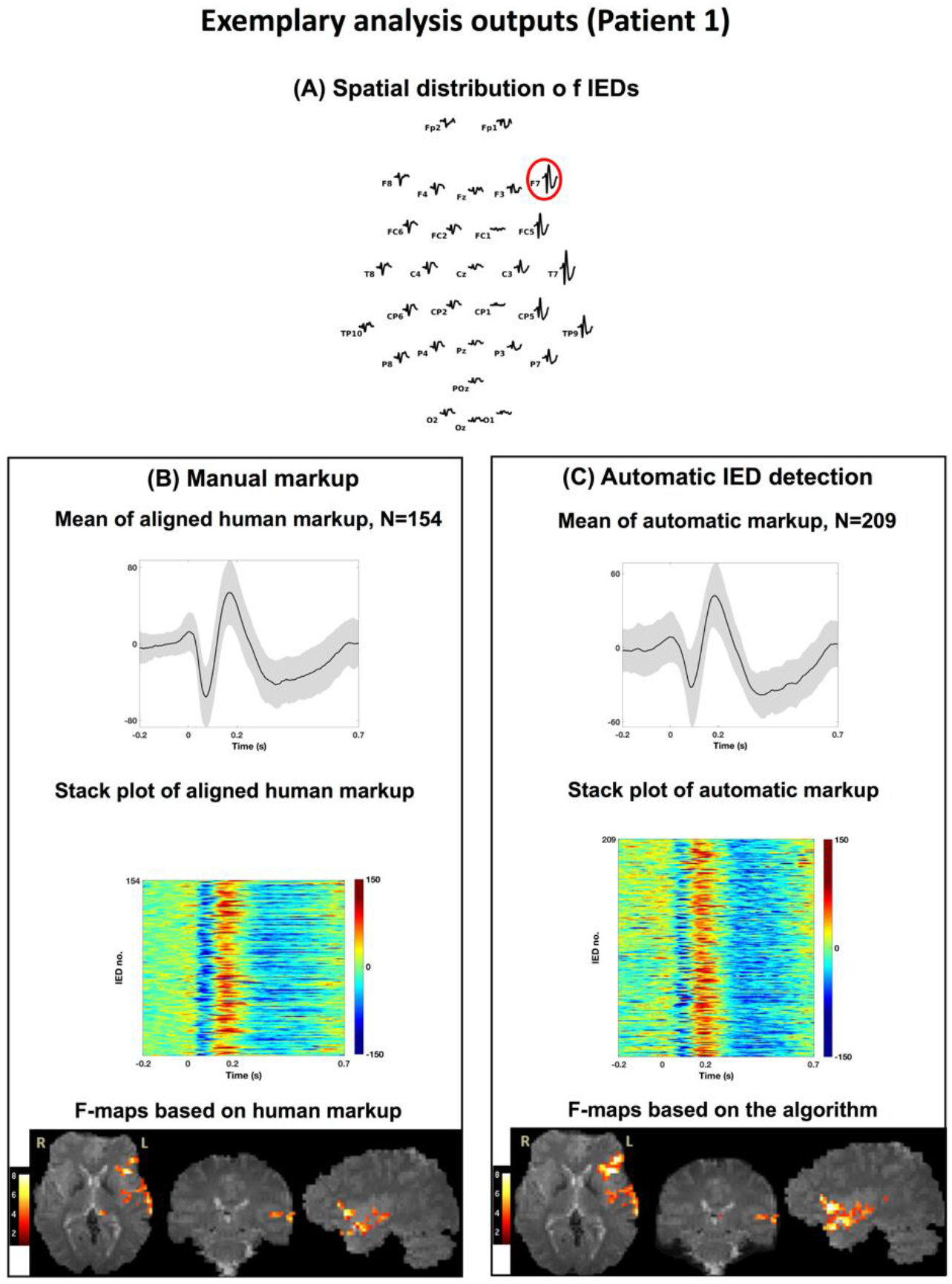
A representative illustration of automatic spike detection (A) and human markup (B) for Patient 1. Top panel shows the spatial distribution of IEDs and the target electrode (F7). Left/right panels show the mean and stack plot of aligned IEDs as well as their associated BOLD inter-subject variability maps. For this dataset, the number of automatically detected spikes was 209 in contrast to 159 manually marked spikes.

**Figure 3:**
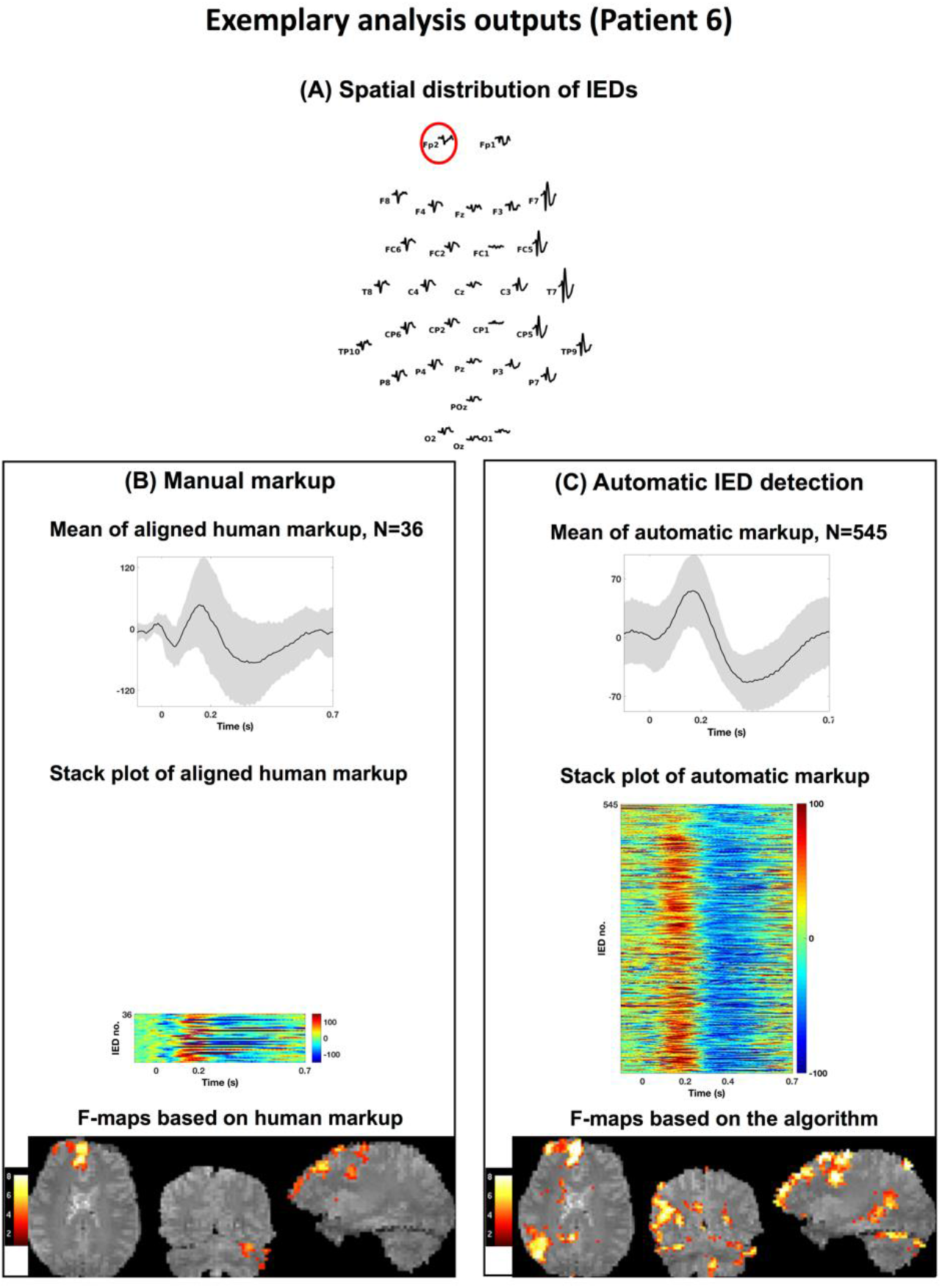
A representative illustration of automatic spike detection (A) and human markup (B) for Patient 6. Top panel shows the spatial distribution of IEDs and the target electrode (Fp2). Left/right panels show the mean and stack plot of aligned IEDs as well as their associated BOLD inter-subject variability maps. For this dataset, the number of automatically detected spikes was 545 in contrast to 36 manually marked spikes.

**Figure 4:**
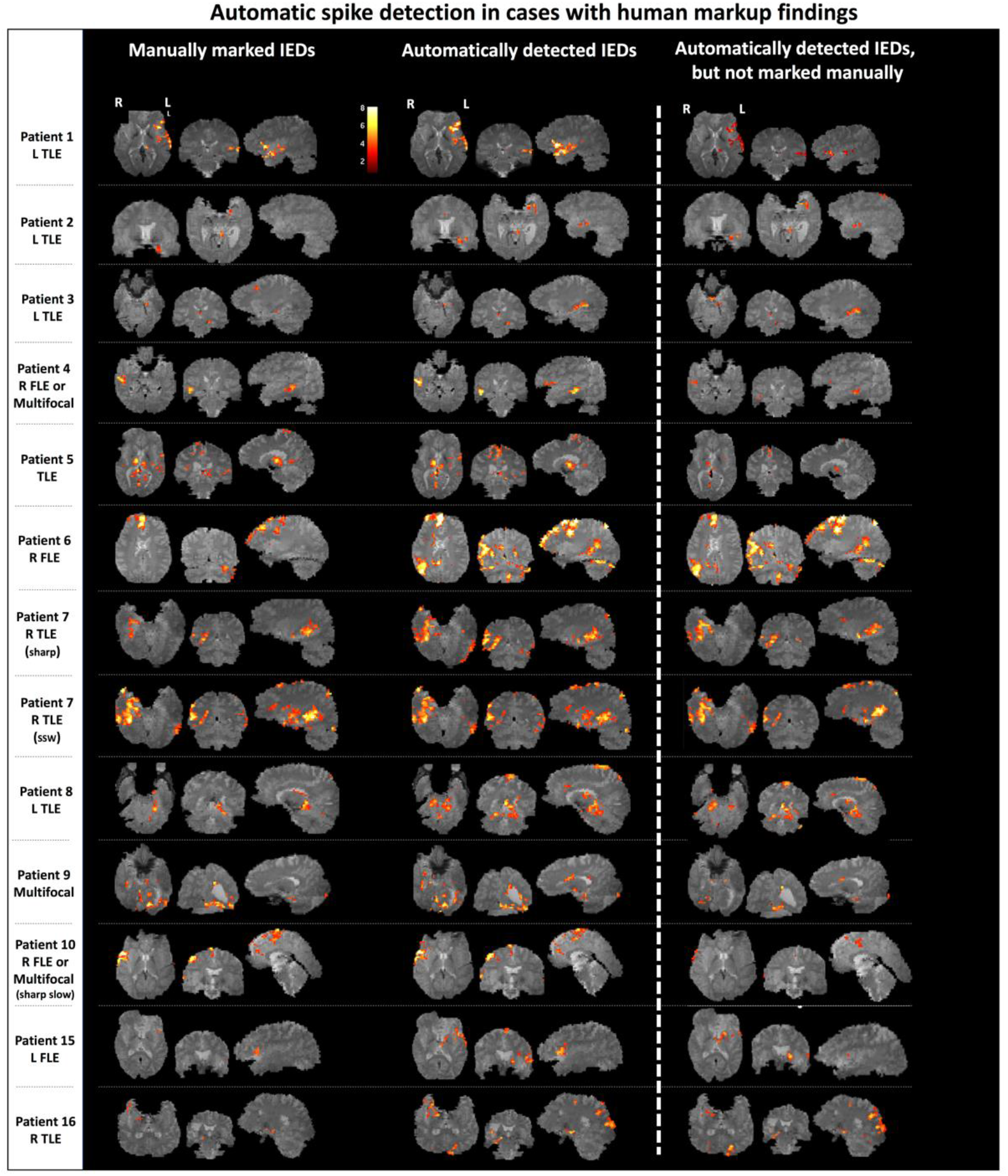
Left panel: BOLD responses based on human markup (left column) and automatic IED detection (right column) of 13 datasets for which at least one markup approach led to the detection of significant BOLD changes in EEG-rsfMRI analysis. Right panel: BOLD responses based solely on manually missed spikes. All T-maps have been thresholded at a voxel-wise significance level of p<0.001 and cluster-wise p<0.05.

The conventional Cohen’s к (Cohen 1960) was used as measures of interrater agreement among the two experts. The к value was interpreted according to the standard guidelines: > 0.81 very good agreement, 0.61-0.80 good, 0.41-0.60 moderate, 0.21-0.40 fair and < 0.20 poor agreement (Altman 1991).

## 3 Results

### 3.1 Automatic IED detection has a short processing time

The most time-consuming and demanding task in any standard EEG-rsfMRI analysis of epilepsy datasets is manually locating interictal epileptiform discharges in EEG recordings by a trained EEG expert. While it would typically take several hours for a human expert to mark epileptogenic events in our one-hour EEG recordings and prepare files with IED onset and duration details, the automatic IED detection method was able to detect IEDs and export an output text file containing IED timings in less than two minutes for all cases (see Table 2 – right column).

### 3.2 Automatic IED detection finds more spikes than human markup

The number of automatically detected IEDs was significantly greater than manually detected IEDs (Wilcoxon rank sum test with p = 0.012). Specifically, the number of manually marked IEDs by the EEG expert ranged from 15 to 587, while the number of automatically detected epileptogenic events varied from 28 to 1,406 (Table 2).

### 3.3 Automatic IED detection improves the statistical power of EEG-rsfMRI F-maps

Human markup led to statistically significant clusters of BOLD changes in only 13 out of 19 EEG-rsfMRI maps (from 16 subjects). On the other hand, statistically significant clusters were present in all 19 EEG-rsfMRI maps using automatic spike detection. Across subjects, the peak F-value across all suprathreshold voxels was higher with automatic spike detection compared to manual markup (Wilcoxon rank sum test, p = 0.031 - see Table 2). Importantly, the number of detected fMRI voxels in the automatic IED detection approach was not significantly correlated with the corresponding threshold values *η_thr_* (Pearson’s r = −0.24 p = 0.32), suggesting that lower ‘similarity’ thresholds (i.e., a more lenient inclusion of ‘automatic spikes’) do not necessarily lead to greater number of suprathreshold voxels in the F-maps.

### 3.4 EEG-rsfMRI maps of automatic and human markup are spatially comparable

Two independent reviews were comparable when evaluated the spatial similarity of EEG-rsfMRI maps between manual and automatic IED markup (expert 1: 6.1 ± 2.4 s.d., expert 2: 6.5 ± 1.9 s.d., see Table 3). Pairwise Cohen’s ͊ coefficients across two sets of review scores for the two types of spatial EEG-rsfMRI maps was 0.759 (very good agreement between the two EEG experts).

#### 3.4.1 Cases in which both human markup and automatic IED detection show significant BOLD clusters

In 13 out of 19 analyses, both human markup and automatic IED detection algorithm elicited statistically significant BOLD changes in EEG-rsfMRI analysis (see Table 3 and Figure 4). In most cases, additional BOLD inter-subject variability identified with automated IED detection were proximate to significant BOLD changes seen on the maps of human markup and consistent with other clinical and imaging evidence (see Table 1, for full clinical information and clinical imaging findings). In two cases (Patient 6 and Patient 8), however, the BOLD response related to automatic IED detection method provided additional information to the EEG-rsfMRI analysis of human markup. Next, we discuss the results of these two patients in more details.

##### Patient 6

This patient was a 26-year-old male whose frequent seizures involved discomfort in his chest, and turning head to the right followed by swearing and repetitive movements. EEG-rsfMRI analysis based on manual IED markup highlighted BOLD changes cross right frontal cortex, midline structures and left cerebellum, suggesting the involvement of a focal frontal epileptogenic network. EEG-rsfMRI analysis of the automatically detected IEDs, on the other hand, showed right parietal lobe inter-subject variability in addition to the previously highlighted brain regions by human markup. This additional region was fully consistent with the origin of patient’s eight stereotypical seizures recorded during EEG-rsfMRI as well as his independent PET results.

##### Patient 8

This patient was a 41-year-old female with seizures characterized by right hand aura of a “crawling feeling” in her hand, followed by a typically generalized seizure. Significant BOLD response based on human markup was detected mainly across left temporal lobe. Based on long-term video EEG monitoring, the patient had interictal discharges with a complex field arising in the left temporal and left posterior quadrant. Her ictal events appeared to have a left posterior quadrant and right hemisphere onset. BOLD changes revealed by the timing of automatically detected IEDs showed that both hemispheres may be involved in epileptogenic networks, a finding which is supported by the video EEG monitoring data.

#### 3.4.2 Cases where human markup led to no EEG-rsfMRI result

In the remaining 6/19 analyses, the IED timing obtained from manual markup did not lead to any significant BOLD changes in EEG-rsfMRI analysis (Figure 5). In most of these cases, IEDs identified by automatic spike detection method revealed significant clusters of BOLD changes which were in line with other available clinical data (PET, SPECT-see Table 1). Next, we review these cases in more details.

**Figure 5:**
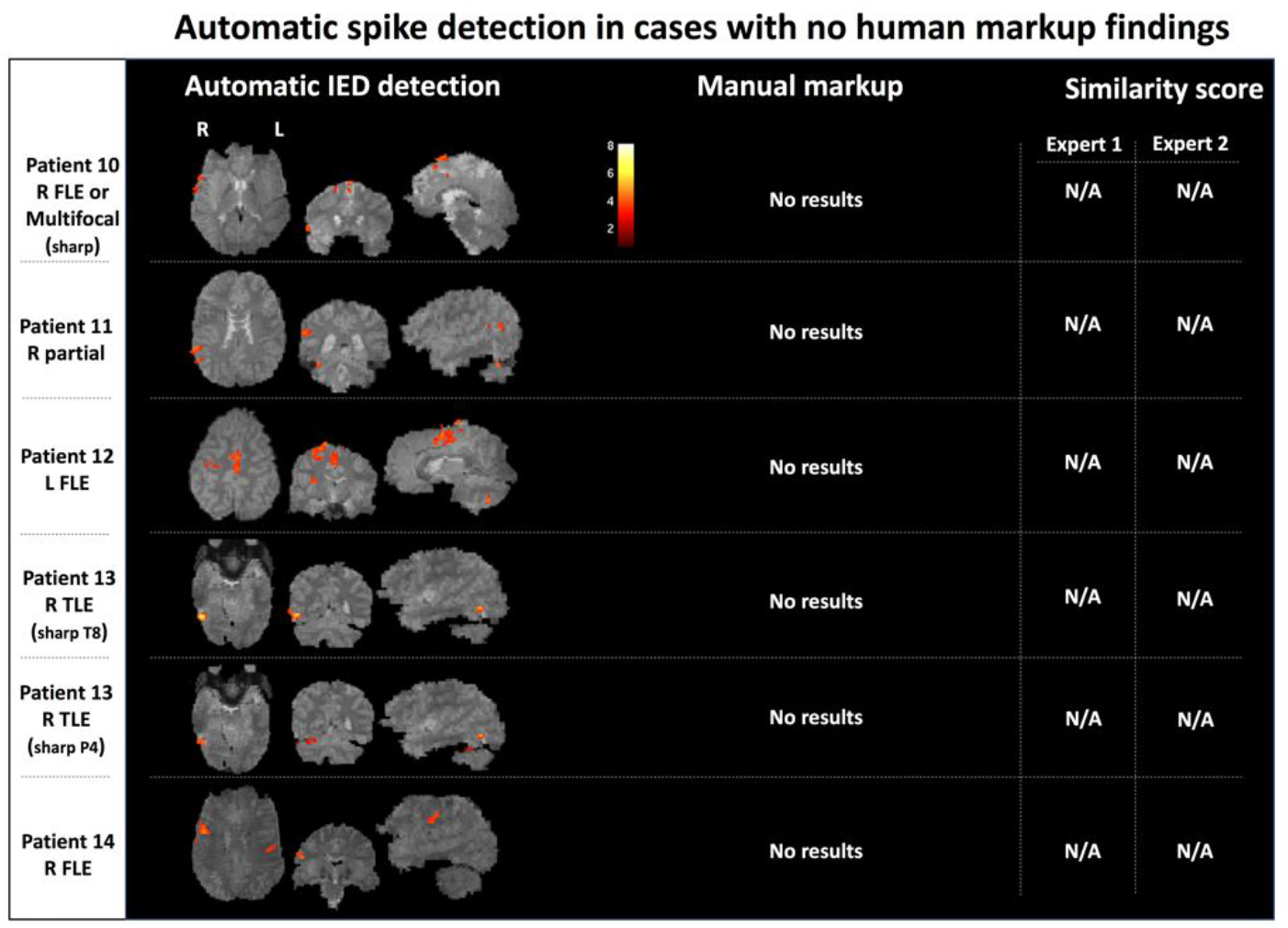
Significant clusters of BOLD changes based on automatic IED detection of five datasets for which manual markup led to no significant results. Therefore, no similarity score by the EEG experts has been given. All F-maps have been thresholded at a voxel-wise significance level of p<0.001 and cluster-wise p<0.05.

##### Patient 10 (sharp IEDs)

This patient, a 14-year-old male, had focal seizures involving clonic jerking down the left side, focal dyscognitive seizures associated with left sided jerking and a psychiatric history with visual hallucinations. His EEG showed bi-frontal epileptiform activity, maximum in the right frontal region at FP2 and F4. Similarly, ictal SPECT showed right frontal changes. EEG-rsfMRI analysis of automatic IED detection method revealed BOLD inter-subject variability in the right frontal regions including midline, middle frontal gyrus, cingulate and temporal lobe. These activated brain regions were consistent with regions identified by nuclear imaging as well as with BOLD signal related to another type of his typical IED included in this study (i.e., sharp-slow - see Table 1).

##### Patient 11

This patient, a 19-year-old female with a 7-year-history of focal seizures beginning in the left hand with some secondarily generalized attacks, had very frequent discharges at C4 on multiple EEG studies. Her MRI showed a possible subtle area of focal thickening involving the depths of the right postcentral sulcus. PET studies revealed hypometabolism in the right post-central region. Stereo EEG confirmed the involvement of the depths of the post-central in epileptogenic focus. The patient subsequently underwent a surgical resection of the suspicious abnormal brain tissue, with significant improvement in her seizure frequency. Interestingly, EEG-rsfMRI based on automatic spike detection method showed significant BOLD changes in right parietal lobe - rather distant from the regions identified by other modalities, but within the same hemisphere.

##### Patient 12

This patient, an 11-year-old female, had a complex left temporal lobe epilepsy or multifocal with possible subtle left parahippocampal dysplasia. She had occasional generalized convulsion seizures, absence seizures characterized by staring periods, with no aura, as well as focal seizures with head/eye deviation to the right. The epileptogenic focus could never be accurately localized. Multiple EEG video-monitoring studies pointed towards left frontal lobe. The patient did not have any nuclear imaging. EEG-rsfMRI based on automatic spike detection method revealed BOLD signal diffuse inter-subject variability in midline and right hemisphere, which is not consistent with other available very limited data.

##### Patient 13

This patient was a 27-year-old female whose weekly seizures began at age 8 and involved impaired consciousness, characterized by blurred vision and were associated with an epigastric sensation, rubbing hands and occasionally lips smacking. Multiple MRI scans showed no lesion, while numerous EEG studies showed focal T8 spikes and ictal SPECT study showed regional right posterior temporal involvement. EEG-rsfMRI analysis based on automatic spike detection method and involving two different types of patient’s characteristic IEDs showed a highly focal area in right inferior temporal posterior brain region, consistent with other clinical findings. Subsequently, intracranial monitoring confirmed this region as the origin of epileptogenic activity. The patient underwent a surgery and is now seizure free after about 24 months of follow-up.

##### Patient 14

This patient was a 25-year-old male whose focal seizures began at the age of 11 years and involved complex visual auras, cognitive and behavioral automatisms, generalized convulsions as well as hemi-clonic activity involving the left side. He also appeared to have some progressive cognitive decline through the 14 years of his seizures. The localization of epileptogenic focus was unclear. His EEG showed interictal frequent discharges in the right anterior quadrant both at F8, F4 and T4 with rare O2 spikes. His anatomical MRI showed diffuse atrophy. Ictal SPECT showed a right mid frontal seizure focus on a background of widespread cortical metabolic abnormality in the right cerebrum, and medial right occipital lobe and right temporal lobe involvement. Automatically detected IEDs in EEG-rsfMRI analysis were associated with significant clusters of BOLD variability in right mid frontal regions, which was consistent with other clinical observations.

## 4 Discussion

In this study, we show that automatic IED detection can reduce the burden of manual EEG markup without compromising data quality. Statistical parametric maps derived from automatic EEG markup had higher statistical power than the ones with manual EEG markup. The additional events seen in automated detection reiterated the pattern of activation seen in the human markups. These results support the use of automatic IED detection for focal epilepsy studies, making EEG-rsfMRI studies more routinely achievable in the investigation of intractable focal epilepsy.

After marking up a sample of the epileptiform events, the subsequent average processing time of IED detection using our matched filtering approach is less than one minute (see Table 2 and Table3). The resulting EEG-rsfMRI F-maps are comparable or better. This contrasts with a typical EEG markup time of several hours by an experienced EEG reporter for a one-hour EEG recording. This approach solves two major issues with EEG fMRI. Firstly, not every event has to be identified individually. Marking every epileptiform event in an hour-long EEG study is difficult, stressful and not the primary skill of the typical reporting EEG individual. Confident identification of a sample of epileptiform events, as needed with our method, is a more typical activity and easily performed. Secondly, the time saving of only marking a few events in the lengthy record makes EEG-rsfMRI a practical step in the clinical evaluation of patients.

Our results also suggest that, even with the best human mark-up, there may be a large number of IEDs throughout an interictal EEG recording which are missed by EEG experts. This is probably because of the uncertainty of recognising low voltage events, Patient 6 is an example of this, where with the automated markup a clearer picture of epileptic networks emerged showing involvement of the association cortex.

Previous studies have suggested a direct relationship between the number of IEDs included in the statistical analysis and the percentage of voxels surviving the statistical thresholding (Flanagan et al., 2009, Gkiatis et al., 2017). Our study confirmed that including more IEDs improves the statistical power of EEG-rsfMRI analysis. The ‘new’ epileptogenic events detected by the automated method enhance the statistical power of the test and revealed (previously hidden) elements of the patient’s epileptogenic networks.

It may have been possible that the automated detection selected ‘noise’ events with characteristics similar to the marked epileptiform events. This is a potential issue that applies to all spike detection techniques as ‘spike-like’ features that match our algorithm may be present in EEG recordings (but actually be noise). However, the third column of Figure 4 shows that, as a group, the new IED’s detected by the algorithm led to biologically meaningful maps of BOLD changes in all patients. Therefore, if there is detection of any additional ‘noise events’ that were not epileptiform it is not a significant problem in our cohort. Separate analysis of only the additional events showed a pattern similar to the original definitely epileptiform events.

Our IED detection algorithm has been validated against human markup. In other words, it gives the same result from marking up just a sample set of discharges as marking up the whole study and all epileptiform events. The brain maps from manual and automatic IED detection were comparable to each other, with additional power in the automated markup. The pattern shown in the additional events ‘marked up’ by the automatic algorithm were similar. This demonstrates that automatic spike detection is a simple, fast and potentially robust tool.

We found it encouraging that in six of our refractory focal epilepsy patients who had no statistically significant results in the standard EEG-rsfMRI analysis with human markup, automatic spike detection provided additional significant fMRI voxels in plausible areas for most of these cases (see patients 10 to 14 - Figure 5). These ‘newly detected’ brain regions were compatible with other available clinical information and imaging modalities such as PET and ictal SPECT (see patient 10, 13 and 14 - Figure 5).

It is important to reinforce that our purpose in this paper was not to evaluate the clinical information provided by the EEG fMRI study, but to assess the automated markup method against human markup. Reliable validation of EEG-rsfMRI results is not a trivial issue, especially when it comes to clinically complicated focal epilepsy patients we report here who often have no clear-cut structural epileptogenic lesion on conventional MRI. In general, surgical resection of the presumed seizure focus with seizure freedom is the gold standard to evaluate this. Our study group included complex patients with active EEGs who mostly did not go on the surgery. As far as we can go (and as can be seen from Figure 4 and 5 as well as the case descriptions), the activated areas are plausible as being linked to the clinical epilepsy phenotype in most cases, including those where activations were only seen when using the automated method.

We consider the current approach as a helper to EEG expert. In essence, the EEG expert only needs to mark a few epileptogenic EEG events to obtain a template definition, and the remainder of the EEG markup will be finalized by the algorithm. Our analyses suggest that automatic spike detection can obtain EEG-rsfMRI results typically in less than a minute by using as few as 15 manually marked spikes (see the fourth column in Table 2). This process is bound to save time and cost of medical staff while providing with the same or improved results as human markup. In order to facilitate the replicability of our findings by other researchers, we are releasing the MATLAB implementation of our IED detection approach available to public (*https://github.com/omidvarnia/Automaticfocalspikedetection*.).

Previous studies have reported good clinical sensitivity using template-based, feature based and topography-based automatic EEG detection methods (Grouiller et al., 2011, Tousseyn et al., 2014, Hao et al., 2018). It is common to use outside-scanner EEG for making a template (in the case of homogenous spikes) or extracting features (in the case of spikes with different temporal and spatial morphologies) and search for similar ‘features’ across in-scanner EEG recordings. Whether an IED template from outside-scanner EEG can be used for inside-scanner EEG dataset with our method has not been tested in this study but might be a future direction for research. Outside-scanner EEG differs from our approach in that the inside-scanner EEG is affected by MRI gradient noise and electrical induction by movement including ballistocardiogram artifacts.

Given that our spike detection method requires a spike template based on human IEDs mark-up, the template matching methods cannot be used for detecting epileptogenic events with inconsistent IED morphology (variable time and shape of epileptic events) such as paroxysmal fast activity. While this may seem to be an issue in patients where spike morphology differs, practically we were able to obtain ‘biologically plausible’ results in our complex cohort despite this. Another limitation is that the spike detection algorithm used in this study only incorporates information of a single target EEG electrode, but the presentation of IEDs may be reflected in a field consisting of multiple EEG channels. A recent study incorporated multiple EEG channels within an automatic IED detection framework. This was done by using a predictive model (deep learning) to include a wider range of IED types in multichannel EEG (Hao et al., 2018). While this is a very elegant solution, it requires lots of data and it is computationally and time demanding. We emphasize that our simple and fast automatic IED detection approach is the ‘polar opposite’ to deep learning algorithm that demands massive amount of data and heavy computational power. We believe that ‘complex’ machine learning techniques and ‘simpler’ signal detection algorithms have complementary roles when it comes to incorporating automatic spike detection as a practical step in epilepsy studies.

## 5 Conclusion

Automatic spike detection is a simple and fast method for reproducing comparable and, in some cases, even superior results in contrast to manual EEG markup in EEG-rsfMRI analysis of epilepsy. Our study shows that automatic detection of interictal epileptiform discharges can be effectively used in epilepsy clinical settings. This work further helps in translating EEG-rsfMRI research into a fast, reliable and easy-to-use clinical tool for epileptologists.

## Acknowledgments

We would like to thank Dr Linda Dalic and Dr Moksh Sethi from Austin Health for the expert review of EEG-rsfMRI statistical parametric maps and Ms. Mira Semmelroch for assistance with patient recruitment and data acquisition.

Author Contributions: AO designed and implemented the spike detection method, conducted the analysis and drafted the manuscript. MK contributed in acquisition and preprocessing of data, conducted the analysis and significantly contributed in writing the manuscript. MP substantially contributed in the discussions and writing the manuscript. GJ provided significant clinical and technical advice, contributed in the discussions and manuscript write-up.

